# Policy driven changes in animal research practices: mapping researchers’ attitudes towards animal-free innovations using the Netherlands as an example

**DOI:** 10.1101/337063

**Authors:** S. Bressers, H. van den Elzen, C. Gräwe, D. van den Oetelaar, P.H.A. Postma, S.K. Schoustra

**Author notes:** Corresponding author (SS). These authors contributed equally to this work.

## Abstract

Reducing the number of animals used in experiments has become a priority for the governments of many countries. For these reductions to occur, animal-free alternatives must be made more available and, crucially, must be embraced by researchers. We conducted an international online survey for academics in the field of animal science (N=367) to explore researchers’ attitudes towards the implementation of animal-free innovations.

Through this survey we address three key questions. The first question is whether scientists who use animals in their research consider governmental goals for animal-free innovations achievable and whether they would support such goals. Secondly, responders were asked to rank the importance of ten roadblocks that could hamper the implementation of animal-free innovations. Finally, responders were asked whether they would migrate (either themselves or their research) if increased animal research regulations in their country of residence restricted their research. While nearly half (40%) of the responders support governmental goals, the majority (71%) of researchers did not consider such goals achievable in their field within the near future. In terms of roadblocks for implementation of animal-free methods, ∼80% of the responders considered ‘reliability’ as important, making it the most highly ranked roadblock. However, all other roadblocks were reported by the majority of responders as somewhat important, suggesting that they must also be considered when addressing animal-free innovations. Importantly, a majority reported that they would consider migration to another country in response to restrictive animal research policy. Thus, governments must consider the risk of researchers migrating to other institutes, states or countries, leading to a ‘brain-drain’ if policies are too strict or suitable animal-free alternatives are not available. Our findings suggest that development and implementation of animal-free innovations are hampered by multiple factors. We outline three pillars concerning education, governmental influence and data sharing, the implementation of which may help to overcome these roadblocks to animal-free innovations.

## Introduction

Animal research has played a critical role in many scientific and medical achievements of the past century. Animal models are used across many fields, including fundamental, biomedical, behavioural, military and agricultural research [1]. Around the world, quality of life has been greatly improved by the research, medicines, treatments and safer environments that have been developed as a consequence of animal-based research in these fields. However, the ethical issues associated with using animals and increased concern regarding animal wellbeing [2], together with concerns regarding the translatability of animal models [3] and practical difficulties of using animals [4], are gaining importance. In line with this, the principles of 3R (Replacement, Reduction and Refinement) described by Russell and Burch [5] have been embedded in national and international legislation and regulations on the use of animals [6, 7]. An example of such international legislation is the EU directive 2010/63/EU, which concerns European wide implementation of the 3R policy [8]. However, the exact success rates of these 3R-related policies towards animal-free innovations is difficult to measure. Evidence from the field suggests that the transition towards animal-free research is moving slowly. For example, funding for studies that use alternative methods is relatively low compared to animal studies [9] and journals that focus on animal-based experiments are generally of higher impact than those that focus on alternative models [10].

Low update of animal-free innovations is partly due to a lack of insight by policy-makers into the preferences and needs of researchers. Additionally, researchers tend to use well-known, widely available methods in their experiments. Furthermore, domestic legislation can easily be bypassed with collaborations abroad, since the research community is a mobile group that often works across institutions, states or countries with varying policies regarding animal research. These and other factors that hamper successful implementation will in this study be referred to as ‘roadblocks’. Given these roadblocks, it is important to investigate the attitude of scientists towards the implementation of animal-free innovations. This is a relatively unexplored terrain in the success of 3R policies. As long as the implementation of animal-free innovations remains limited and researchers remain unaware of alternatives, such methods will not gain traction within scientific disciplines. Therefore, attempts to improve the implementation have been made.

A recent example of a governmental policy advocating the implementation of animal-free innovations is a goal set by the Dutch government, which aims to become the world-leading country in animal-free innovations by 2025 [11]. This will be addressed as the ‘2025-goal’ in the remainder of this article. Questions that arise from such a goal include whether researchers would be supportive, and whether they think this goal would be practical and achievable. In addition, to promote the communication between governmental instances and academia, we tried to gain insight into the most important roadblocks of the implementation of animal-free innovations. In this study, researchers from both the Netherlands and other countries were asked to comment on these questions. A future consequence of restricted legislation concerning animal research may be migration of researchers to other areas with less strict regulations, reducing the country’s competitiveness in research [12]. By investigating the probability of researchers migrating to other institutes, states or countries because of stricter legislations, the consequences of such governmental measures can be estimated.

Given the global increase in concern for animal welfare, it is likely that other governments will set similar goals regarding the use of animal-free innovations in research. Mapping the attitudes of both Dutch and foreign researchers towards the Dutch 2025-goal provided insights from those who are subject to the goal, as well as outside perspectives. In this matter, we can map the attitude of the Dutch researchers, but also that of others who might experience similar goals in their own country. Furthermore, the hypothesis that a proportion of researchers may move to another location in response to more strict regulations was addressed. Evidence of stagnation in knowledge-development as a consequence of forced restricting in legislation will be presented and discussed. Gaining insights into the above will allow for exploration of the success of governmental policies and the attitude of researchers towards the implementation of animal-free innovations.

## Methods

An international online survey asked participants about the 2025-goal, a list of potential roadblocks, and their willingness to migrate as result of governmental influences on the implementation of animal-free innovations. Data management, security, and integrity of the survey was approved by the Social Sciences Ethics Committee at Radboud University in Nijmegen, the Netherlands (registration number: ECSW2017-3001-466), and was endorsed by rector Prof. Han van Krieken, license holder for animal research at Radboud University and Radboudumc.

### Sample selection

Scientists at academic centres in the regions of Nijmegen, Rotterdam, Utrecht, Amsterdam, Maastricht and Groningen, as well as large academic centres located in the United States and in countries surrounding the Netherlands were invited via email to participate in the survey. The same process was used for companies in the Netherlands that perform animal-based research. All potential participants were currently working in or had worked in any field related to animal experimentation, alternatives or policy. The informed consent, that was sent along with the survey, included a request for participants to share the survey with others who might be interested. Therefore, the non-participation rate for this study is unknown. In total, 457 participants responded to the survey, but only responders working in the academic sector (N=382) were selected for analysis as they form a uniform and comparable group, Fig 1. Additionally, students working in an academic setting (N=17) were excluded because they are still relatively new in this field. This resulted in a study population of 367 researchers.

**Fig 1.**
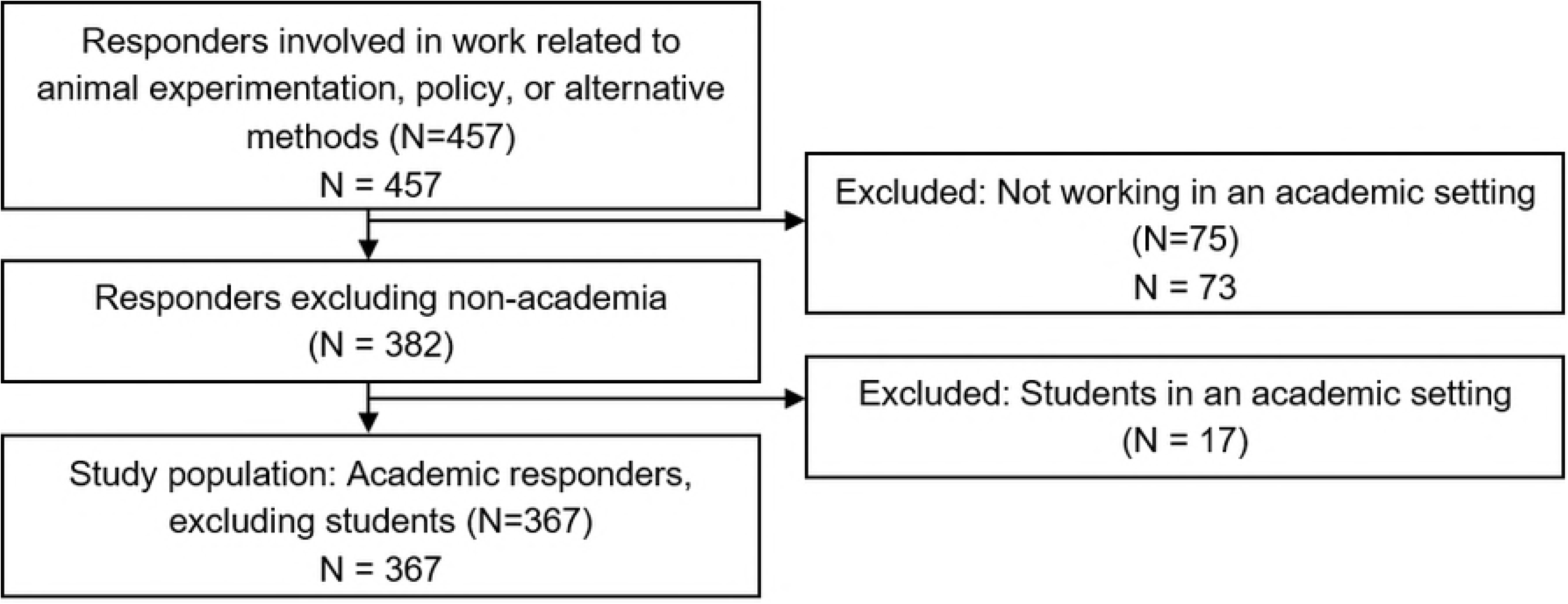
Selection of study population concluding a group of 367 researchers working in an academic setting.

### Survey procedure and measures

The survey was available from March 18 to March 27, 2017. To obtain an overview of the opinions and thoughts about the 2025-goal, responders were briefly introduced to the 2025-goal. Hereafter, the following questions were asked: *“What is your opinion about the number of animals currently used in experimentation in the field you are working in?”, “Should research be animal-free?”, “Is the 2025-goal achievable?”*, and *“Would you support the 2025-goal?”*. After this, responders were asked to rate the potential importance of a set of ten potential roadblocks. These roadblocks were identified from previous studies conducted in the Netherlands that addressed possible issues regarding the implementation of alternatives for biomedical sciences [13, 14]. The resulting list was narrowed down to ten roadblocks that were used to establish a ranking based on the outcome of the survey. After each question, responders had the opportunity to elaborate on their answers in free text boxes. After this, demographic information was gathered, including information about educational background, nationality, and whether the participant was currently working with animals. Based on their answer to the latter question, responders were directed to specific questions regarding animal models and alternatives, and were asked for their personal opinion about different statements, including the question whether researchers would consider moving to another country due to changes in regulation regarding animal experimentation.

### Statistical analysis

Age was expressed as mean with standard deviation (±SD). Categorical variables were expressed as absolute numbers and percentages. Comparisons between subgroups were carried out using chi-squared tests in SPSS statistics 21. The answers provided via the free text boxes were analysed manually and summarized to obtain an overview of the perspectives shared by scientists working in academia.

## Results

The population investigated in this survey included scientists working at universities or research centres, and employees of companies in any field related to animal experimentation, alternatives or policy-making within the field of animal research. A total of 457 responses were obtained, of which 367 researchers in an academic setting were selected. Academic researchers were in our opinion the best choice to select our date on because they form the biggest and most influential group of people involved in the execution of animal experimentation.

Of the sample as a whole, the mean age was 38 (±11) and roughly equal numbers of men and women responded, 56% and 44% respectively. Regarding the level of education attained, the distribution of the responders was as follows: 38% PhD and/or MSc, 25% Principal Investigator, 16% post-doc, 21% other, Table S1. Of all responders, 74% were directly involved in animal research at the time of the survey. More detailed information on the general demographics can be found in Table 1, and Tables S1A-S1D in S1 Text.

**Table 1.**
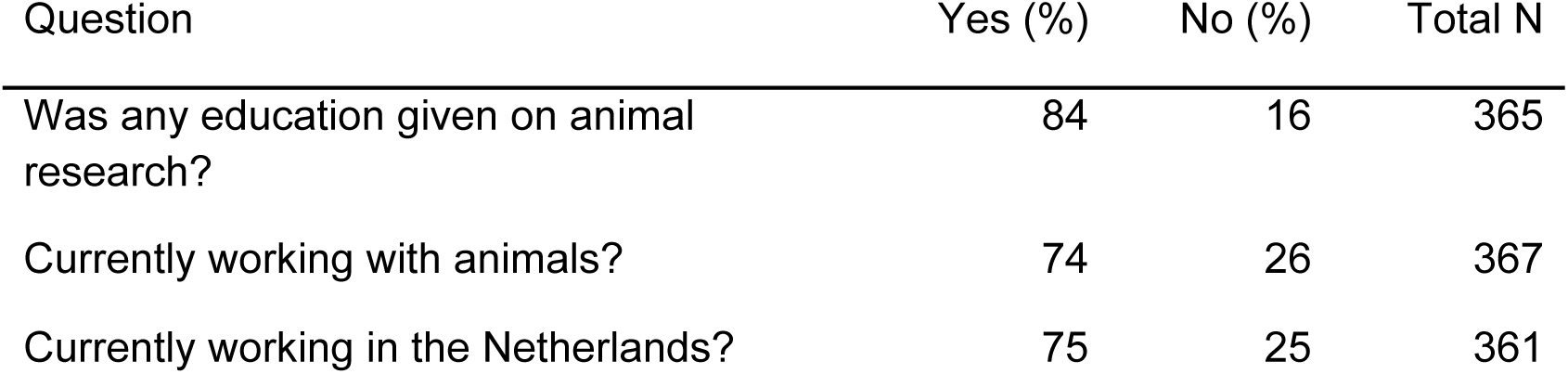
General demographics of survey respondents (N=367)

| Question | Yes (%) | No (%) | Total N |
| --- | --- | --- | --- |
| Was any education given on animal research? | 84 | 16 | 365 |
| Currently working with animals? | 74 | 26 | 367 |
| Currently working in the Netherlands? | 75 | 25 | 361 |

### Achievability and support towards the 2025-goal

As the 2025-goal is a recent example of governmental influence on the use of animals and the stimulation of animal-free innovations, this setting was used to map the attitude of researchers towards such a goal and, according to their perspective, rank the importance of the selected roadblocks. Furthermore, the influence of such a goal on migration of researchers or their research was investigated.

By studying the preferences and needs of researchers regarding the achievability of the 2025-goal and their support towards this goal, a more successful implementation of animal-free innovations could be achieved. Among researchers, animal studies and its regulations are a delicate but lively topic. 85% of the responders expressed themselves by using one or multiple free text boxes to substantiate their opinion about this topic. Of these responders, 43% made use of every free text box, indicating the close involvement of researchers with this topic.

The majority of the researchers (71%) shared the opinion that the implementation of the 2025-goal is not achievable in their field of research. However, 40% of the responders indicated that they would support such a goal, Fig 2A. Many believed that, at this moment, knowledge on alternative methods is not sufficient to abandon the use of animals completely. Nevertheless, researchers expressed that if the government would invest heavily in alternatives for animal models, the goal should be possible eventually. However, they did not expect significant change on such a short timescale as 2025.

Because the quite prevalent difference between how researchers responded to the question towards the achievability of the goal and to the question whether they would support the 2025-goal, we further split up the analysis by comparing several groups. The opinions of researchers working with animals versus those who do not were compared, Fig 2B and Fig 2C. 78% of the researchers working with animals share the opinion that the 2025-goal is not achievable, compared to 53% of the researchers who do not work with animals (*p ≤ 0.001)*. In addition, 36% of the former group would support the 2025-goal, comparing to 54% of the latter group (*p ≤ 0.01)*, Fig 2D and Fig 2E. Comparing researchers working in the Netherlands versus those who do not show no statistical differences in both achievability and supportiveness to the 2025-goal that was used as an example.

**Fig 2.**
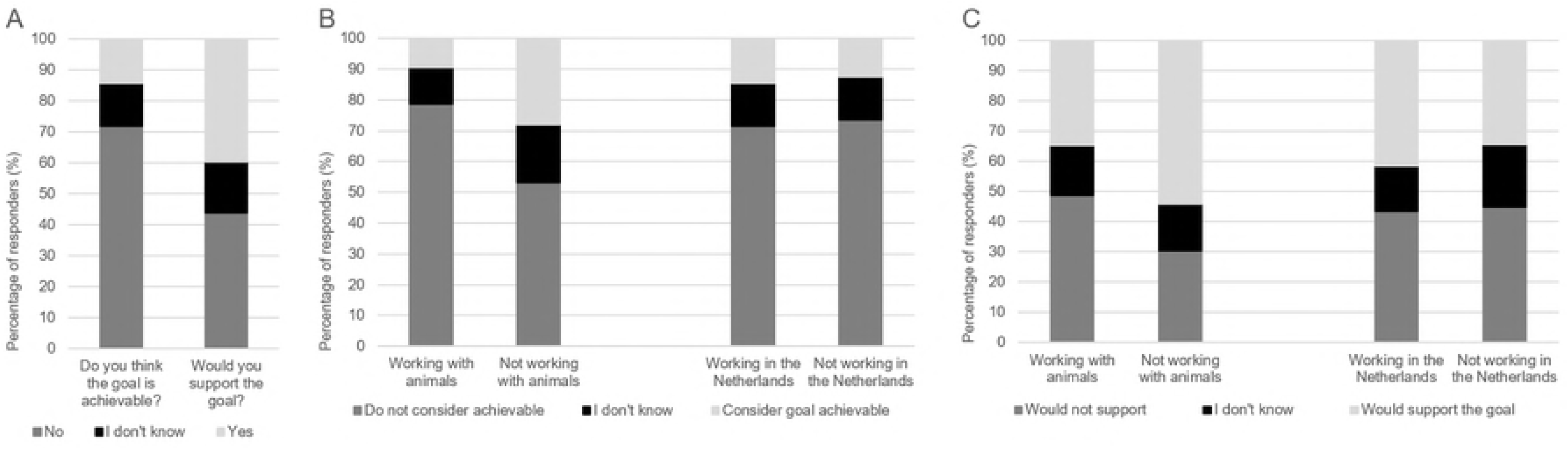
Overview of opinions towards the 2025-goal according to researchers participating in the current study: (A) Response for the questions whether the responders thought the 2025-goal is achievable and whether they would support it (N=367). (B) Achievability of the 2025-goal divided into researchers working with animals (N=271) and researchers working without animals (N=96), together with the division of researchers working inside the Netherlands (N=280) or outside of the Netherlands (N=87). (C) Supportiveness of the 2025-goal divided into researchers working with animals (N=271) and researchers working without animals (N=96), together with the division of researchers working inside the Netherlands (N=280) or outside of the Netherlands (N=87).

### Ranking the roadblocks

In order to indicate the flaws in communication between governmental instances and researchers, we gained insight into the most important roadblocks of the implementation of a goal like the 2025-goal, as seen by researchers. Ten roadblocks were pre-selected based on previous literature, Table 2. Definitions of the selected roadblocks can be found in S2.

**Table 2.** Ten most frequently listed roadblocks for implementation and development of alternatives.

|  |  |  |  |
| --- | --- | --- | --- |
| I | Alternatives are not animal-free | VI | Pressure to conform |
| II | Awareness is lacking | VII | Publishing in high-impact journals |
| III | Costs of implementation | VIII | Reliability |
| IV | Differences in regulation | IX | Research funding |
| V | Ethical issues | X | Time/effort to develop alternatives |

In total, 64% of the responders ranked the roadblock ‘reliability’ as ‘very important’, Fig 3. To put that score into perspective, the roadblock with the second highest percentage of the category ‘very important’ is ‘time/effort to develop alternatives’. This roadblock was ranked as ‘very important’ by 29% of the responders. When we combine the scores of ‘very important’ and ‘important’, only the aforementioned roadblocks have a majority of responders giving these scores. Even though differences exist between the ranking of ‘reliability’ and ‘differences in regulations’ or ‘ethical issues’, all roadblocks were ranked as (very) important to some extent.

**Fig 3.**
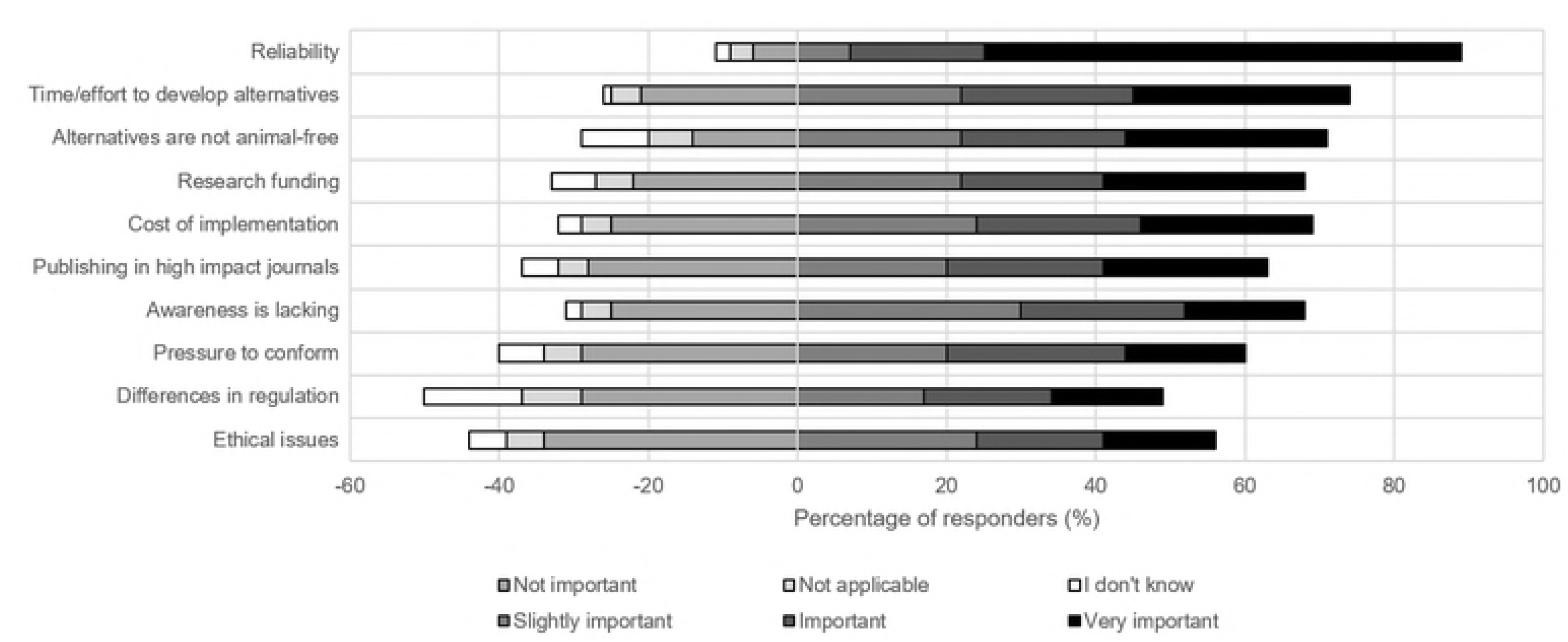
Overview of roadblocks ranked according to their importance as stated by the researchers: The importance of the different roadblocks was scored using six categories: ‘very important’, ‘important’, ‘slightly important’, ‘not important’, ‘I don’t know’, and ‘not applicable’.

### Migration of researchers

A potential and rather serious impact of regulations such as the 2025-goal could be that researchers feel forced to leave institutes, states or countries in order to keep their research going. To further investigate the possibility of scientists leaving and their opinion on migration due to regulations, responders working with animals were directed to further in-depth questions concerning animal research. Responders were asked whether they would consider moving to another place if their animal research were no longer allowed where they were currently working. Of the responders, 46% would consider moving themselves or their research, 23% answered maybe, and 31% would not, Fig 4A. However, the responders who would not or would maybe consider moving frequently gave as additional motivation that they would collaborate with research institutes abroad, rather than moving themselves. This means that (a part) of their research will be moved abroad after all. Overall, more than half of the responders would consider to move either themselves or their research.

To determine a possible relation between the age of responders and their willingness to migrate when their research were no longer allowed, answers were compared between different age groups, Fig 4B. Responders with an age between 30 and 39 had the highest percentage of researchers who would consider to migrate. The highest amount of responders who were uncertain, were people of age 20 till 29. Responders with an age between 50 and 59 had the highest percentage of responders who would not consider moving.

To determine whether responders who are already working abroad would consider migration more easily than researchers who are working in their native country, the answers of these two groups were compared with each other, Fig 4C. Of the researchers working abroad, 67% would consider moving, compared to 41% of the responders working in their native country. The percentage of researchers who were uncertain was similar in both groups. A slightly bigger percentage of researchers working in their native country answered that they would not consider moving (24%), compared to responders working abroad (20%).

**Fig 4.**
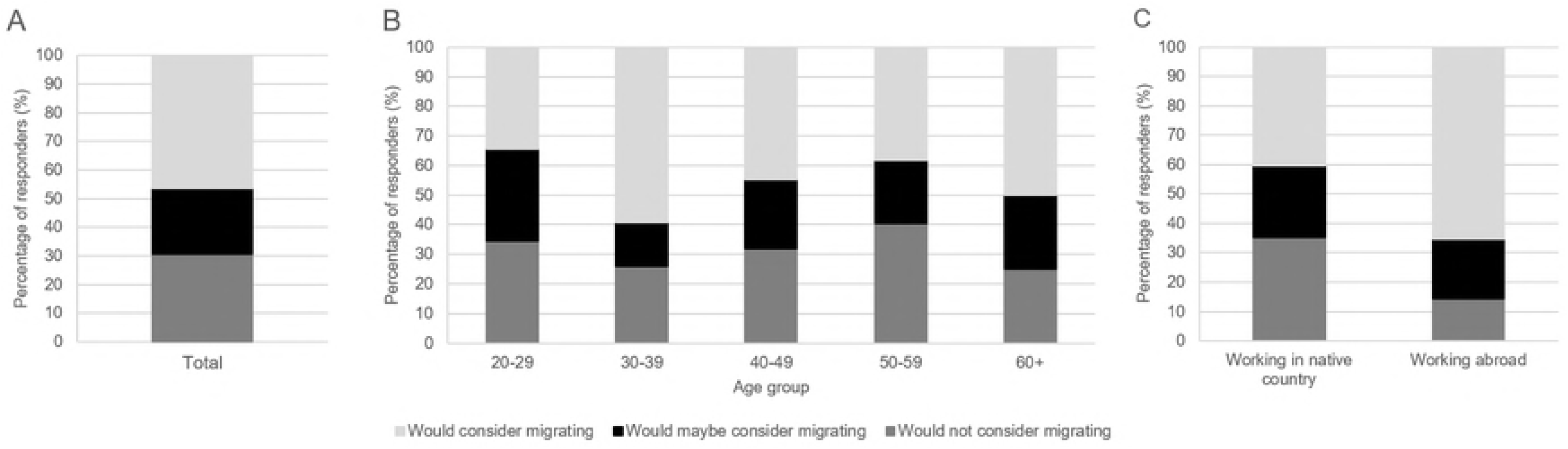
Willingness to migrate due to governmental legislation: (A) General opinion on the question whether the responders would consider to migrate due to stricter governmental legislation (N=271). (B) Willingness to migrate divided by age groups of 20-29 (N=58), 30-39 (N=81), 40-49 (N=47), 50-59 (N=47) and 60+ (N=12) years old. (C) Willingness to migrate comparing researchers working in their native country (N=193) and working abroad (N=49).

## Discussion

In this paper, we elucidated the thoughts and opinions of researchers concerning support and achievability of governmental goals to stimulate innovations in animal-free research. We were able to determine the most important roadblocks in the implementation of animal-free methods, as seen by researchers. Finally, the paper demonstrated that researchers are more willing to migrate as a result of stricter legislation.

### Implementation of governmental goals

Whereas approximately half of the responders would support governmental regulations concerning implementation of animal-free innovations, 71% of the researchers share the opinion that implementation of the 2025-goal is not yet achievable in their field of science. Implementation of innovations that focus specifically on reducing and/or replacing animal models is not simple given the complexity of animal research and its purposes [14]. Therefore, investment of governmental agencies across the world in the refinement of necessary animal experiments might result in minimizing stress and discomfort amongst animals used for experimentation. Furthermore, readily available innovations could be used more efficiently and researchers should be made (more) aware of them. Besides that, cross-sectoral and multidisciplinary cooperation could be stimulated in order to improve innovative developments towards alternative methods for animal research [14]. With this cooperation, new developments can be shared across scientific or national borders.

### The roadblocks in perspective

All the roadblocks for the implementation of animal-free innovations included in the survey were ranked at least ‘slightly important’ and the total of responses stating ‘important’ were more than the total of ‘not important’, ‘not applicable’ or ‘I do not know’. This implicates that all roadblocks could be considered to be at least of some importance. Therefore, a multidisciplinary approach is advisable as a solution. Given the partial similarities of solutions to separate roadblocks, we consolidated these solutions in three main pillars: education, government and data sharing. In Table 3, a full list is presented of all roadblocks and what pillars may form the solution to tackle these roadblocks, as indicated by a checked (black) or unchecked (white) box.

**Table 3.** Relation of the roadblocks towards the three pillars. Checked boxes indicate that the given pillar would suit the needs to tackle the given roadblock.

| Roadblock | Education | Government | Data sharing |
| --- | --- | --- | --- |
| Alternatives are not animal-free | ☐ | ☐ | ☒ |
| Awareness is lacking | ☒ | ☒ | ☒ |
| Costs of implementation | ☐ | ☐ | ☒ |
| Differences in regulation | ☒ | ☒ | ☒ |
| Ethical issues | ☒ | ☒ | ☐ |
| Pressure to conform | ☒ | ☐ | ☐ |
| Publishing in high-impact journals | ☐ | ☐ | ☒ |
| Reliability | ☒ | ☒ | ☒ |
| Research funding | ☐ | ☒ | ☐ |
| Time/effort to develop alternatives | ☒ | ☒ | ☒ |

The *education*-pillar includes universities, which could provide their students and employees with better training and access to knowledge regarding alternatives and their development. Institutions could offer courses on the development and implementation of alternative methods in order to educate their employees. This makes students and employees more aware of the opportunities of animal-free experimentation and may promote the choice to consider alternatives in the future.

The second pillar is the *government*, which can influence the implementation of animal-free innovations in multiple ways. First, funding is required since large amounts of money will be required in order to stimulate the development and implementation of new animal-free innovations. Documentation of the available animal-free methods should also be centralized in a reliable open access database in order to increase awareness and usage of the existing alternatives. Finally, for a smooth transition of legislations or goals like the 2025-goal, it is required that all stakeholders are aware of their responsibilities and expectations of others. Hence, a proper communication between the government and the public has to be established.

The pillar on *data sharing* includes accessibility of ‘work-in-progress-data’ that allows researchers to obtain more insights in what is going on in their field. Researchers tend to not share research data prior to publication. However, when these data are not shared, other researchers remain unaware that someone is already working on a certain topic. This might result in unnecessary duplication of experiments. In addition, publishing negative data is not incentivized, often being rejected outright from journals or only accepted in low-impact journals. This makes it of low priority for researchers. This however leads to duplication of findings since other researchers remain unaware of these negative results and may therefore perform the same experiments just to conclude the same negative results. Unnecessary repetition of experiments must thus be prevented and can be solved by an increase in data sharing. This will result in a lower amount of sacrificed animals and will prevent waste of valuable time and resources.

### Policy driven migration of researchers

Van Noorden reviewed the global migration of scientists and the factors that play a role in this process [12]. As cited from the article, the goal was to “identify underlying trends in scientists’ movements, investigate what is driving them and explore how they may change” [12]. The majority of our responders would consider moving to another institute, state or country when their research were no longer allowed in the country where they were currently working. Similar to our results, Van Noorden presented that an ‘authoritarian political system and restricted freedom’ were seen as barriers for emigration to that country by 93% of the responders [12]. Factors that were seen as incentives by the majority of his responders included: ‘improved quality of life’ (88%), ‘more research funding’ (84%) and ‘better salary’ (77%) [12]. Governmental influence is, therefore, not the only factor, but it does affect considerations of researchers regarding migration or collaboration with other institutes. A higher percentage of responders willing to migrate was expected to be found in the younger age groups, considering Van Noorden’s results [12], as younger people might be more flexible and therefore less tied down to a specific location. Additionally, work-related migration was studied more in depth as people who migrated before might easier migrate a second time than those who are still working in their native country. Both hypotheses were supported by our results, indicating that researchers form a mobile community, willing to migrate or move their research if necessary.

Researchers did express their concerns about the position of the Netherlands and its developments in animal-free innovations compared to other countries. These concerns include that stricter regulations could lead to a drainage of animal research to other countries, which eventually could harm the research climate in the Netherlands. To prevent this, researchers that responded to the survey proposed internationalization of a goal like the 2025-goal. When more countries promote the development of alternative models towards animal research, the risk of negative effects on individual scientific positions could be reduced [14].

Despite the relatively large study population, participation bias could be a limitation of the current study, as the non-respondent rate remains unknown. Although we distributed our survey both in and outside the Netherlands, almost 75% of the responders were working in the Netherlands. This might have affected the answers of our responders, as the Dutch 2025-goal was used as an introductory background. A more international public might have been reached by emphasizing the generality of the 2025-goal to a broader extent. Besides that, explanation of the individual roadblocks was lacking in the survey. Therefore, interpretations of the stated roadblocks might have differed amongst participants. Furthermore, potential important roadblocks might have been excluded from the pre-selected ones. However, as none of the responders mentioned novel roadblocks, it can be concluded that at least the most important roadblocks were included in the survey.

## Conclusion

The 3R principle is becoming increasingly more prominent in legislation concerning animal research. As a result, greater stress is placed on the use and development of animal-free innovations, as is reflected in the 2025-goal of the Dutch government that served as an example in this paper. However, less was known about the attitude of researchers concerning animal-free innovations. This paper demonstrated that researchers take legislation concerning animal research into account, and would consider to migrate when they could not perform their research due to stricter legislation. Hence, if one aims to make a systematic impact in animal research, animal regulation should be coordinated at an international level. If not, research will simply be transferred to less-regulated countries. In addition, researchers clearly expressed their preference that animal-free research should be at least as reliable as the rival animal model. Education, governmental influence and data sharing are tools to optimize the implementation of alternative methods. Ultimately, a structural solution is only possible if animal-free research becomes more appealing to researchers, and not by forcing the community.

## Acknowledgements

The authors thank Dr. Amanda Tilot and Dr. Sonja Vernes for excellent supervision throughout the process, as well as Prof. Dr. Merel Ritskes-Hoitinga, Dr. Matthijs Kox, and Prof. Thomas Korff, for providing essential insights. In addition, we would like to thank the Radboud Honours Academy, in particular Noortje ter Berg, for the support we received and for all opportunities that were given to us while working in this interdisciplinary think tank. Without their help we would not have had the chance to do this research. Lastly, we want to thank Prof. Han van Krieken for his endorsement and all survey participants and anyone else who was somehow involved in this project.

